# Allosteric communication induced by GTP binding sets off a closed-to-open transition in a bacterial dynamin-like protein

**DOI:** 10.1101/2023.01.16.524228

**Authors:** Wibke Schumann, Birgit Strodel

## Abstract

Dynamin superfamily proteins are mechanochemical GTPases that operate in highly oligomeric and highly cooperative superstructures to deform lipid membranes. It is known from the structures of a bacterial dynamin-like protein (BDLP) that binding of GTP and association of BDLP with lipids causes a transition from closed to open hinge 1 that affects oligomerization. We trace this radical, large-scale conformational change at the atomic level with unbiased, replica exchange, and umbrella sampling molecular dynamics simulations. We decipher how GTP loading from the GTPase domain to the distal stalk end is mediated by an allosteric network of salt bridges that act in response to GTP binding and subsequent conformational changes in GTPase domain motifs. Two previously undiscovered motifs have been identified whose movements free the paddle from the GTPase domain, allowing large-scale domain rearrangements. In addition, a novel wide-open state of BDLP reminiscent of human dynamin 1 is discovered. Our results explain several aspects of the BDLP cycle and have broad implications for other members of the dynamin family.

## 1 Introduction

Dynamin-superfamily proteins (DSPs) are mechanochemical enzymes, involved in critical cellular functions like endocytosis, cell division and immune response.^1–4^ Next to flagellar proteins, they generate some of the highest torques known for proteins, in the range of a thousand piconewton-nanometers (or a few attojoule).^1,5^ DSPs differ from the smaller Ras-like GTPases in their lack of accessory proteins, lower substrate affinity, and higher basal hydrolysis rate, which is highly stimulated in oligomers.^6–9^ The oligomers form ordered lattices, rings, or helices, which tubulate membranes.^2^ The conserved GTPase domain of DSPs spans about 300 residues and features an internal GTPase-activating domain to replace the external activation factors.^9–11^ GTP binding leads to oligomerization and stronger membrane association, while GTP hydrolysis leads to fission and oligomer dissociation.^3,12^ GTPase activity needs to be highly concerted, local and fast, but not simultaneous, rather propagating along the supramolecular helix to avoid weak points.^13,14^ In polymerized DSPs, 2–4 GTPs can be hydrolyzed per second, and the open and closed states have lifetimes in the range of a few seconds.^15,16^ The GTP dissociation rate lies in the range of 10-100/s and thus, dimer lifetime of DSPs is short (in the range of hundreds of milliseconds). ^17^ Assembly of a microscopically visible dynamin sheath on a membrane can take up to an hour,^17^ while membrane fission can occur as soon as 10–20 subunits are assembled, with the final fission happening in seconds to minutes.^17^ Several mechanisms for dynamin action have been proposed, among them the poppase, pinchase, and twistase mechanisms. ^18–20^ The poppase action works by extending the membrane stalk, thinning the bilayer, which then ruptures. In the pinchase theory, it is a conformational change that leads to a reduction of the DSP helix diameter and thus to membrane constriction. The twistase mechanism achieves the same effect by forming supramolecular winches, which decrease the number of proteins per turn. Permanent membrane binding ability is neccessary for fusion DSPs, while several cycles of constriction are needed for fission.

Bacterial dynamin-like proteins have been associated with several membrane-related functions, including membrane vesicle formation and membrane fusion. The current model is that bacterial dynamin-like proteins are recruited to sites where homotypic membrane fusion is required. The bacterial dynamin-like protein from *Nostoc punctiforme* (called BDLP henceforth) is supposed to be involved in fusion rather than fission. BDLP has a canonical G domain separated by hinge2 from the neck and trunk region, which are connected via hinge1 Figure 1. The membrane interaction is mediated by a paddle domain. BDLP localizes to the outer leaflet of the inner plasma membrane and could be the cyanobacterial ancestor of the thylakoid-reshaping fuzzy-onion-like protein in higher plants. Membrane binding of BDLP is connected to the dimerization of its G domain, self-assembly of the C-terminal GTPase effector domain, and the paddle region contacting the lipids promoting membrane curvature. This can only be achieved in the open or extended state of BDLP, whose structure was resolved to 9 Å with cryogenic electron microscopy as a dimer loaded with a GTP analogue (GMPPNP, guanosine-5’-[(*β,γ*)-imido]triphosphate) and with the paddles inserted into a lipid membrane in a pincushion-fashion.^21^ This requires a large-scale conformational change from the closed state of BDLP, whose GDP-bound structure was resolved with X-ray diffraction to a resolution of 3 Å. It features an acute triangular arrangement of the G domain, neck and trunk^22^ and is less likely to bind to membranes.^23^ The BDLP is the only DSP for which high-quality structures exist in both the open and closed conformation.

**Figure 1:**
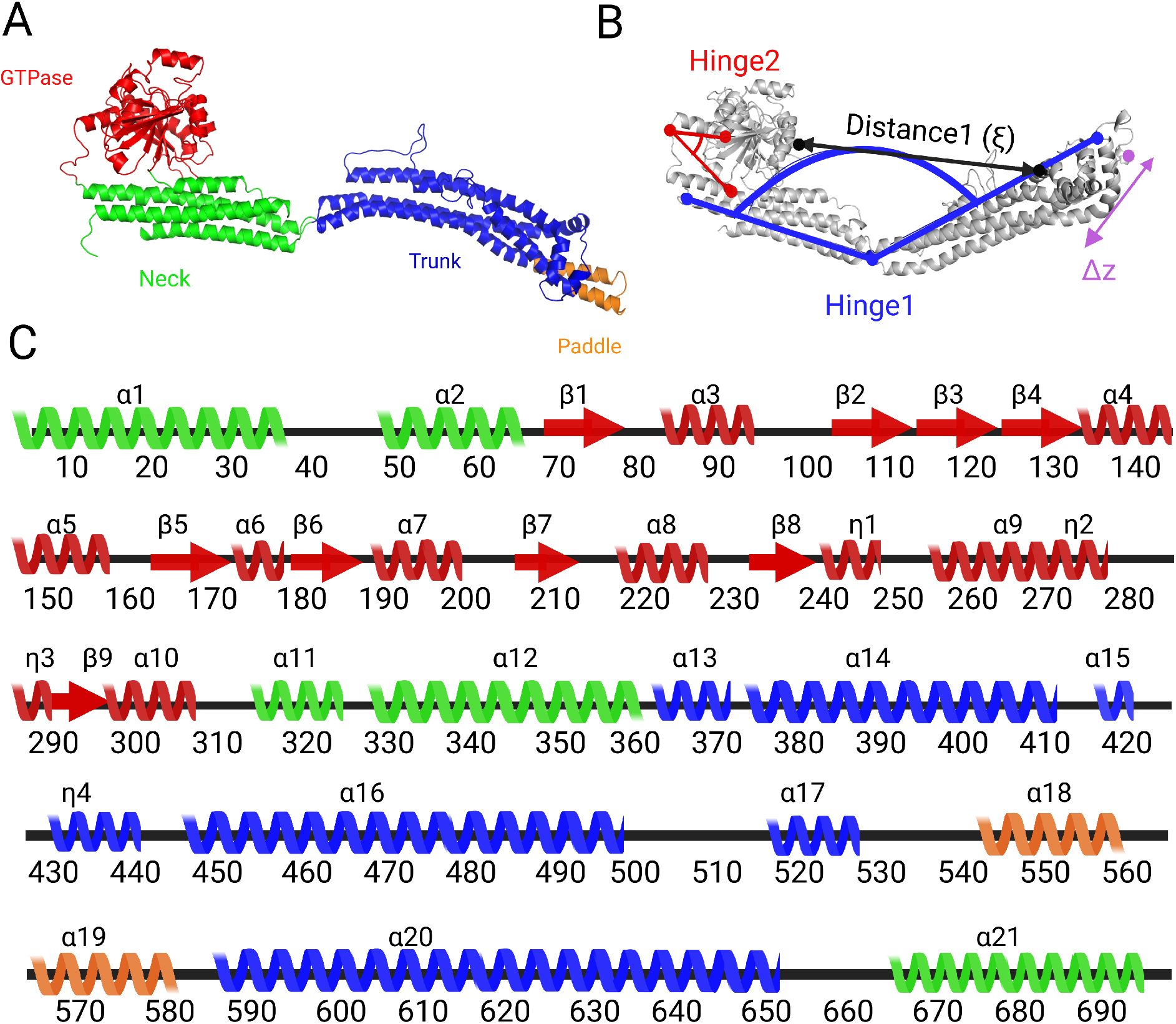
Structural characteristics of BDLP. **A** The open form of BDLP is shown as cartoon and colored red for the G domain, green for the neck, blue for th trunk, and the mmembrane-binding paddle is shown in orange. **B** The open and closed states and their interconversion are characterized by several order parameters. Hinge1 is described by the angle *α* between the C_*α*_ atoms of residues 4, 359, and 587 (blue), with *α* ≈ 175^*°*^ representing the open BDLP conformation and *α* ≈ 40^*°*^ the closed one. Hinge2, which was previously proposed to be an important switch for the GTP loading state, is characterized by the angle *β* between the C_*α*_ atoms of residues 291, 303 and 323 (red). The amount of opening or closing is measured by the distance *ξ* between the C_*α*_ atoms of residues 224 and 453 (black arrow), and the lateral stalk trunk motions are defined by changes along the *z* coordinate (magenta arrow). **C** The secondary structure of BDLP along with the labeling of its helices and *β*-strands is shown.

The aim of the current work is to unravel the molecular details of the closed-to-open transition and provide a rationale for the structural prerequsitites and implications of that radical conformational change. To reach that goal, we perform all-atom molecular dynamics (MD) simulations, which has become an accepted method to fill the gaps left by experimental methods as it provides a higher spatial and temporal resolution than the experiments.^24–26^ Thus far, no all-atom MD simulations on DSPs have been reported yet; only coarse-grained simulations with non-atomic resolution of DSPs are published to date.^27–32^ In our lab, we simulated different guanylate-binding proteins (GBPs), which belong to the dynamin-related superfamily, at atomic resolution, which revealed a large-scale hinge movement that may correspond to hinge1 in the DSPs.^33,34^ Support for such a hinge movement is provided by experimental observation.^15,35^ Nonetheless, the hinge movement sampled in our simulations of GBPs did not involve a complete open-to-closed transition, as suggested by the BDLP structures. We therefore set out to close this gap, using the open and closed BDLP structures as input, to characterize the complete hinge movement for the first time. We employ standard MD simulations, Hamilitonian replica exchange and umbrella sampling MD simulations (HREMD and USMD respectively), and explore the allosteric effects triggered by GTP binding.

## 2 Results

### 2.1 Open BDLP is highly dynamic and can adopt a semi-closed conformation

We started the study by exploring the overall stabilities and local flexibilities of apo- and holo-BDLP in the open and closed states in unbiased MD simulations. The analysis of protein flexibilities, as measured by the root mean square fluctuations of the C_*α*_ atoms (RMSF), revealed that on the simulated timescale of 600 ns, both apo- and holo-BDLP in the closed conformation are stable, while the open forms displayed more flexibilities, yet more so in the case of apo-BDLP (Figure 2). Indeed, the open conformation of apo-BDLP started to spontaneously close (Figure 3A), reaching a semi-closed state with *α ≈* 100^*°*^. In the closed holo-BDLP state, GTP binding induced flexibilities, yet it remained closed. Nonetheless, the different flexibilities in closed apo- and holo-BDLP indicate that GTP loading may have the potential to drive the protein towards the open conformation, while apo-BDLP prefers closed conformations.

**Figure 2:**
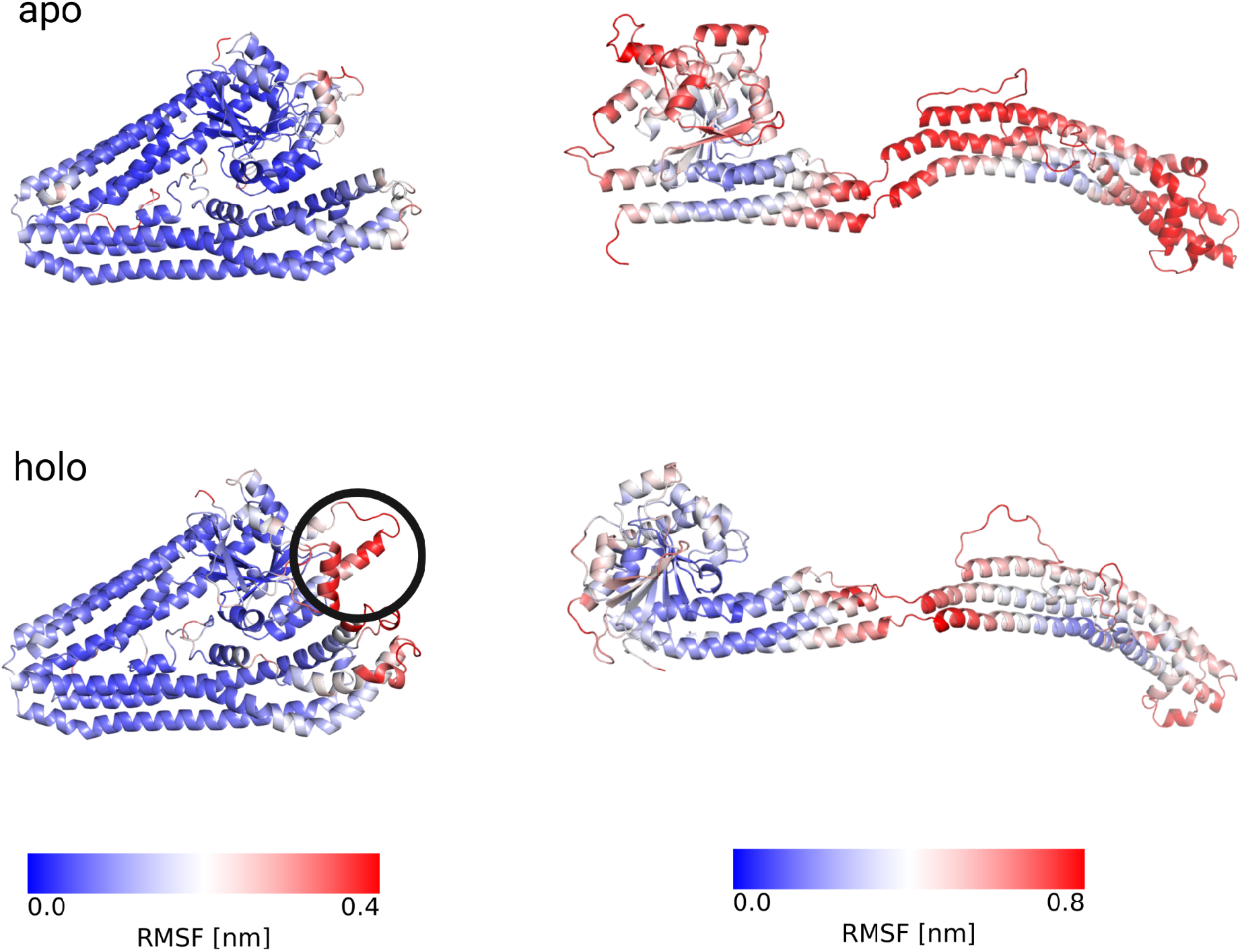
Root mean square fluctuations of the C_*α*_ atoms observed during 600 ns MD simulations of BDLP in different states: apo/closed, apo/open, holo/closed, holo/open. The RMSF values are project onto the cartoon presentations of BDLP, using colors ranging from blue for low RMSF to red for high RMSF. The exact color scales are given below the less mobile closed states (left) and the more flexible open states (right). The simulations revealed the existence of two important structural elements in the G domain that interact with the paddle in the closed form and respond to GTP binding by conformational changes. Because of their flapping motion, we called them flap1 and flap2. Their location is highlighted by a black circle in the representation of the closed holo-BDLP.

**Figure 3:**
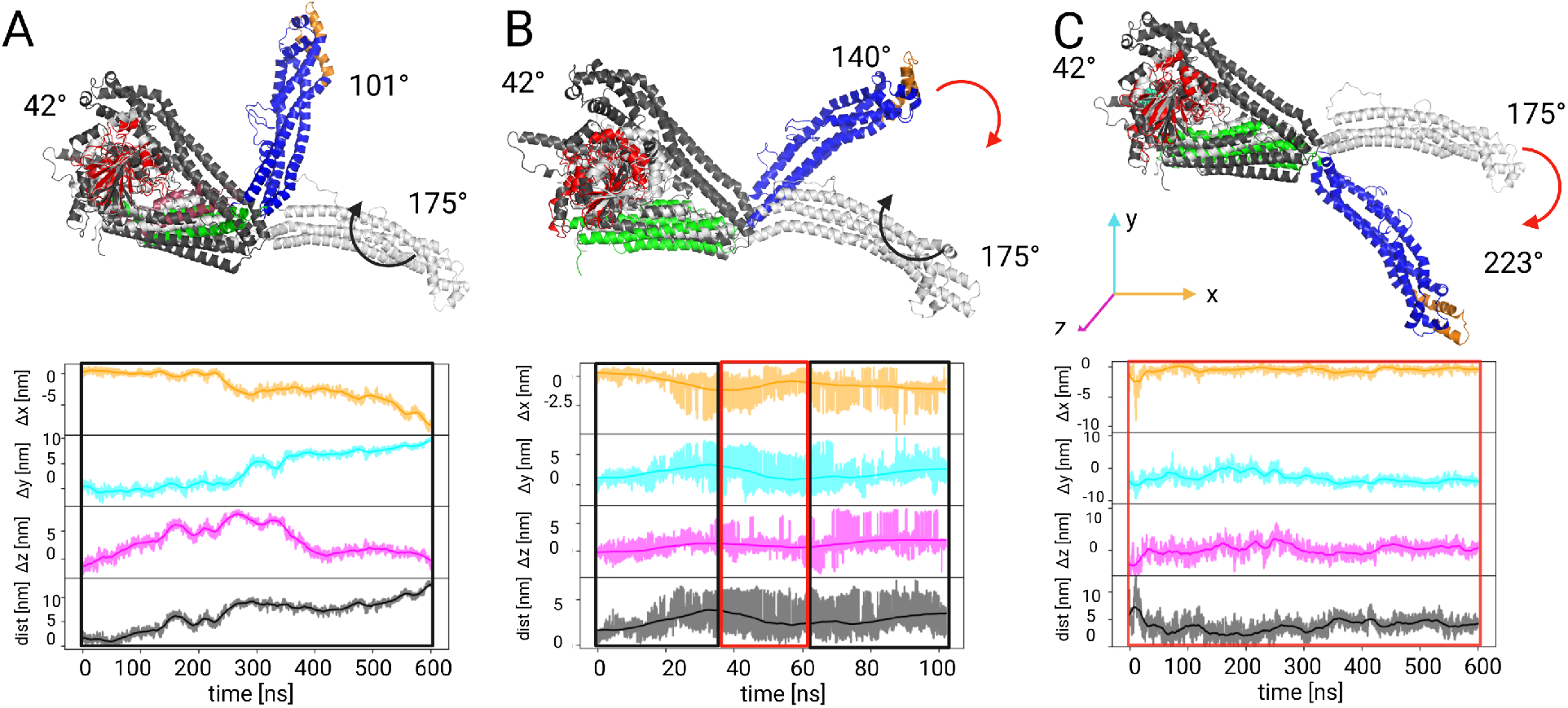
Hinge movement of BDLP illustrated by structures (top) and deviations of the stalk-tip position from its initial location (bottom). The movements are quantified by Δ*x*, Δ*y*, Δ*z*, and overall distance, where the opening and closing directions are indicated by black and red boxes, respectively in the bottom panels. **A** In the MD simulation that started from the open conformation (white) of apo-BDLP, the protein assumed a semi-closed structure with *α* ≈ 100^*°*^. For comparison, the fully closed state is shown (black), which remained closed in an MD simulation that started from this conformation. **B** In the HREMD simulation of apo-BDLP that started from the open conformation, both closing and opening movements were observed. with a mimimal *α* 140^*°*^ that was reached. **C** In the HREMD simulation of holo-BDLP, starting from the open conformation, the conformation not only remained in the open state, but even adopted a wide-open conformation with *α* reaching up to *≈* 225^*°*^.

To further explore possible conformational transitions, an HREMD simulation of apo-BDLP starting from the open conformation was performed. Here, the switching between the open and semi-closed state (here with *α ≈* 140^*°*^) was sampled more often and reversibly (Figure 3B). We then simulated both the open and closed form of holo-BDLP using HREMD. The setup for the closed state confirmed the observation from the MD simulations that there are structural instabilities at the interface between the G domain and the paddle (Figure 2). They result from motions in the G domain set off by GTP binding, especially involving *α*5 (residues 143–156) and *α*9 plus the preceding loop (residues 251–271), which we denote flap1 and flap2 because of their swinging motions further discussed below. The open holo-BDLP, on the other hand, quickly assumed *α* angles over 180°, hereafter referred to as wide-open conformation (Figure 3C), which is highly reminiscent of human dynamin 1 and has not been observed for BDLP before. Another interesting observation is that the open-to-closed motion in BDLP involves movements of the trunk into all three spatial directions, as resolved by the analysis of the stalk tip motions (Δ*x*, Δ*y*, and Δ*z* in Figure 3), so that in fact hinge1 is rather a ball joint, at least a restricted ball joint, than just a hinge as its name suggests.

### 2.2 Hinge1 is not just a hinge but a restricted ball joint

A more general overview of the conformational landscape of BDLP is provided by free energy surfaces. The free energy (Δ*G*) is projected onto different combinations of the order parameters *α* and Δ*z*, which were already discussed above, as well as the distance *ξ* to measure the amount of opening and the angle *β* to assess hinge2 movements. The results in Figure 4A and B are shown for the HREMD simulations of the open forms of apo- and holo-BDLP, respectively. The Δ*G*(*ξ, α*) values for apo-BDLP confirm that it closes spontaneously to reach the semi-closed state, which leads to reductions in both *ξ* and *α* values. However, for angles between 110 and 150^*°*^, different distances between the stalk tip and the G domain can be assumed (around 6 nm and 8–10 nm). This degeneracy indicates that motions in another direction must occur in parallel, and this third dimension is the lateral motion Δ*z*. The projection Δ*G*(*ξ*, Δ*z*) shows that the trunk of BDLP upon closing first moves into positive Δ*z* direction (see Figure 3 for the definition of the coordinate system) and then starts to move into the opposite direction at *ξ ≈* 9 nm. This confirms the conclusion made above that hinge1 is more than a hinge, it can rather be considered a restricted ball joint. The hinge1 movements are correlated with hinge2 movements, as revealed by snapshots in Figure S1. The monomeric BDLP in open conformation was constructed from a dimer structure, where the G domain is rotated against the neck precisely at hinge2, and in part this difference is conserved during simulation. In the closed conformation, the hinge2 helical region was resolved, while the rotated open conformation had to be modeled with a disordered loop region. However, the disordered hinge2 region becomes more helical with GTP being bound.

**Figure 4:**
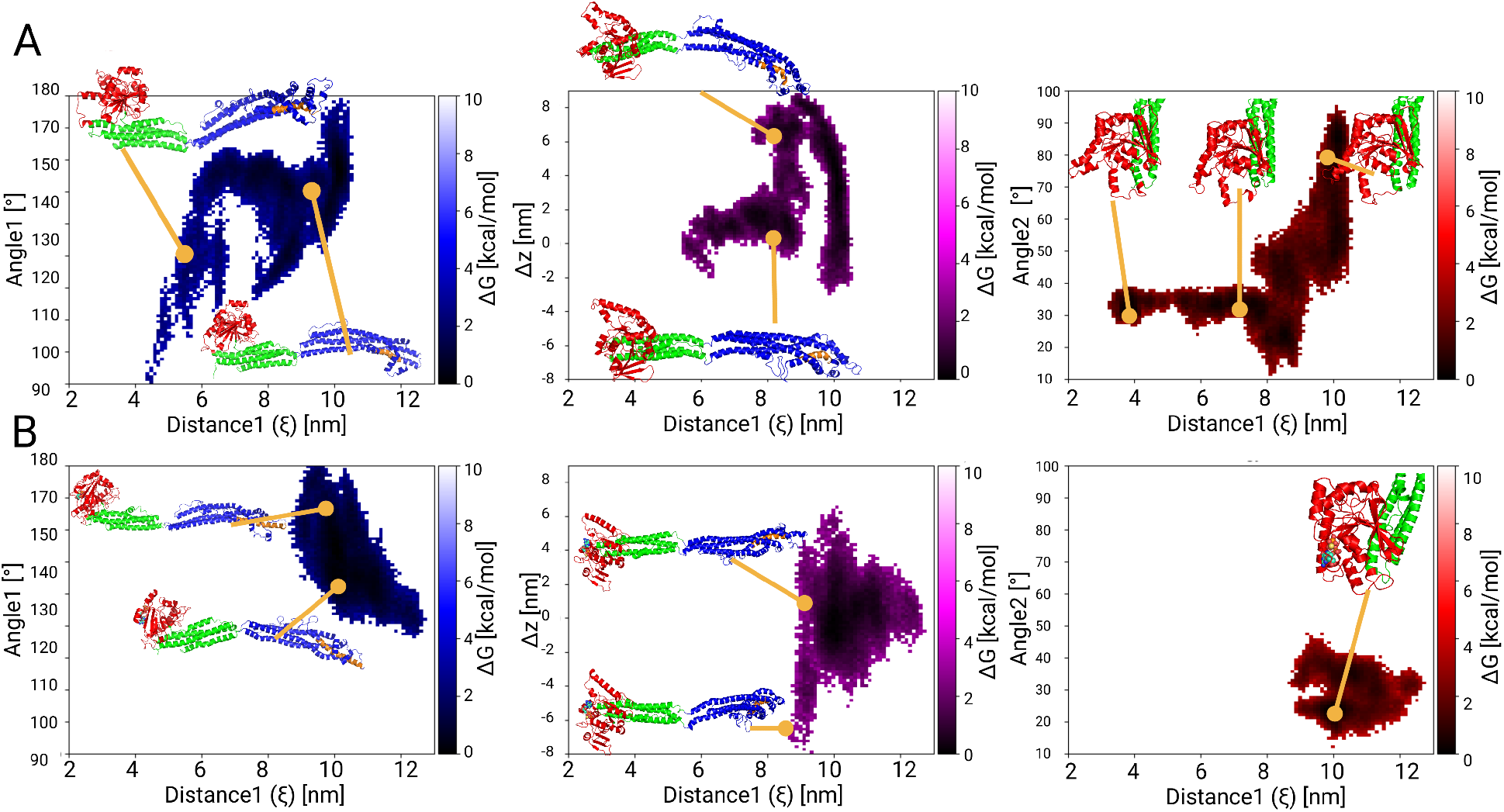
Free energy surfaces as a function of selected order parameters. The free energy is shown as 2D projections Δ*G*(*ξ, α*) (left), Δ*G*(*ξ*, Δ*z*) (middle), and Δ*G*(*ξ, β*) (right), along with representative structures for **A** apo-BDLP and **B** holo-BDLP. Δ*G* values are color-coded according to the scales on the right of the plots. Low Δ*G* values indicate stable structures.

The corresponding Δ*G* plots for holo-BDLP in Figure 4B confirm the observation made above that GTP binding stabilizes the open conformation and even induces a wide-open conformation with *α >* 180^*°*^ and *ξ >* 10 nm. The movements toward the wide-open structure also involve lateral motions of the trunk, yet into negative *z* direction. Thus, using the BDLP presentation as shown here, with the G domain on the left, one can state that apo-BDLP closes with the trunk moving towards the reader, while holo-BDLP wildly opens away from the reader. Since no major closing in holo-BDLP took place, hinge2 did not change much. However, the hinge2 region displays a more structured head-neck conformation, which in apo-BDLP was only reached upon hinge1 closing. Especially at *ξ ≈* 8 nm a coupling between the stalk and the G domain in apo-BDLP must occur, as at that distance not only the lateral movement of the stalk tip abruptly reverts back to Δ*z ≈* 0, but also *β* converts to smaller angles here (Figure S1). The lateral motion restriction could be due to the interaction of a loop in the trunk region (490–510) with the neck, guiding the swivel motion into a more linear approach towards the G domain.

### 2.3 GTP binding sets off long-distance communication

In order to understand the coupling between the G domain and stalk, we continue by analyzing the structural flexibilities of the G domain and how they are affected by GTP binding. To this end, a principal component analysis of the most mobile regions of the G domain was performed, which revealed the swinging motions of the previously mentioned flap1 and flap2 (Figure 5). Especially the closed conformation of holo-BDLP displays pronounced flap1 and flap2 movements to cover GTP. G domains are typically characterized by the presence of certain motifs that are directly or indirectly involved in GTP binding. The only canonical GTPase motif that can be clearly identified in BDLP is the P-loop at residues 76–84. The other motifs deviate, but can be mapped to G2 (or switch I, 102–103), G3 (or switch II, 180–184), G4 (238–241), and G5 (245–268). Moreover, hinge2 also belongs to the G domain (Figure 5) and its conformational change is of relevance for the coupling between the domains.

**Figure 5:**
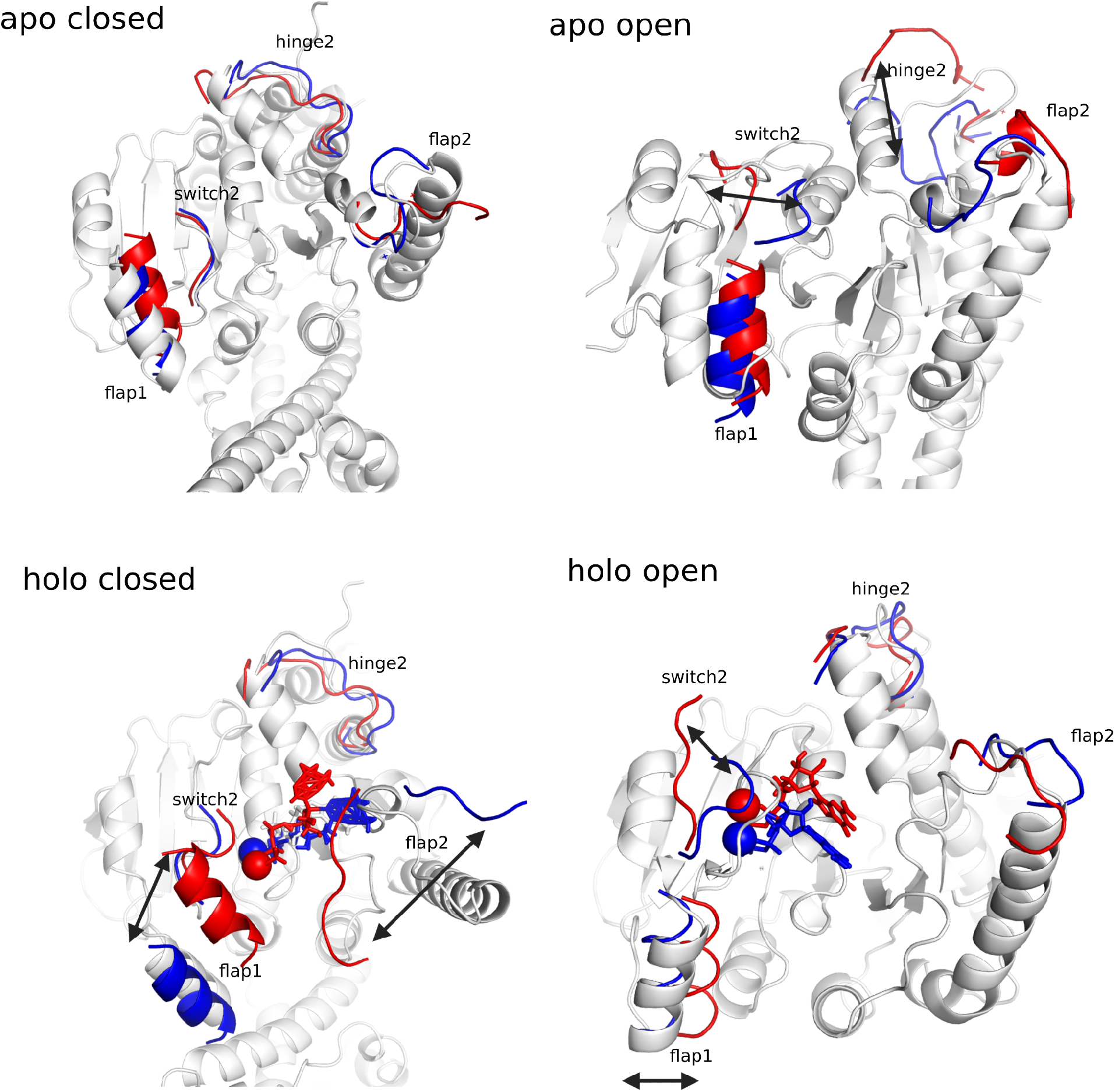
Principal motions of important structural elements of the G domain in apo-BDLP (top) and holo-BDLP (bottom) in the closed (left) and open (right) conformations. Red and blue mark the maximum displacement in either direction of the motions of flap1, flap2, hinge2, and switch2. In the case of holo-BDLP, also the relocations of Mg^2+^ and GTP are shown. The arrows indicate the respective movement.

As previously observed in our studies of murine guanylate-binding protein 2, ^34^ GTP binding has the capability to stiffen the G domain. This also occurs in BDLP when it is in the open conformation. Conformational clusteing of all G domain motifs produced 103 rather ordered clusters for holo-BDLP, while this number sextuples for apo-BDLP. Here, 624 clusters are found and a high variance among the cluster conformations of hinge2 and flap2 is present. However, the situation reverses for the closed form of BDLP. In that case, in the GTP-bound state even more clusters are found than for apo-BDLP: 178 clusters for holo-BDLP with a high variance in hinge2 and flap2 and only 61 clusters with a high degree of structural order for apo-BDLP. This suggests that the open conformation can better accommodate GTP, while GTP binding induces structural instabilities in the closed state that might induce further structural changes in the stalk region.

With the aim to identify communication pathways between the G domain and the stalk, we identified salt bridges that change their occupancies upon GTP binding in the MD simulations of closed BDLP. The most obvious candidate that might function as an allosteric switch is a salt bridge connecting flap1 with the trunk, K154–E438, which is dissolved by a flap1 relocation. In the GTP-bound state, flap1 and thus K154 moved towards GTP and thus away from E438 (Figure 6A), located adjacent to the paddle domain. Two further salt bridges between the G domain and the trunk broke: R221–D454 and R226–E464. Of note is also the salt bridge E348–K502 connecting the neck with the trunk, whose stability also decreased followed GTP binding. Interestingly, these residue pairs are all on the same side of BDLP (which is the right side in the protein presentation in Figure 6B). Contrariwise, on the other side of the protein, there are three salt bridges that gained in strength upon GTP binding: E188–K446 connecting the G domain with the trunk, R352–E645 between the neck and the trunk, and K653–E657 associating the loop region between the trunk and neck with the paddle domain. This weakening and strengthing of salt bridges across the protein strikingly reveals how GTP binding gives rise to conformational changes in the closed BDLP that are expected to facilitate the closed-to-open transition. The salt bridges thus represent important allosteric switches that enable the information flow from the GTP-binding site in the G domain to the different parts of the stalk region. Moreover, the two-sided distribution of weakened/dissolved and strenghtened/newly formed salt bridges further confirms the observation made above that movements of hinge1 are more than a swinging motion but also involve lateral (or shearing) movements. This is further supported by a normal mode analysis of the closed conformations (Figure 7), which shows that in holo-BDLP the two longitudinal sides of the stalk move in opposite directions. This is not the case in the closed apo-BDLP.

**Figure 6:**
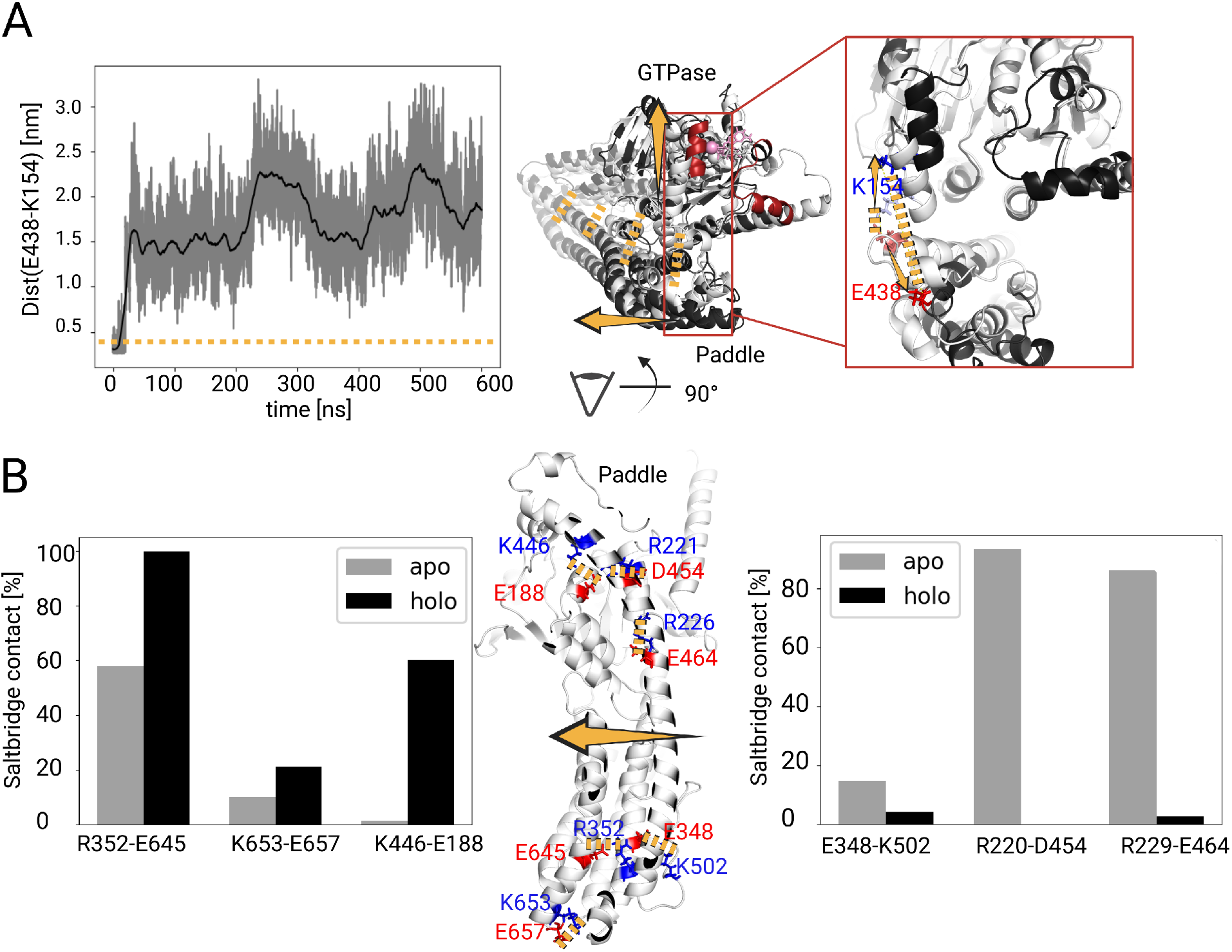
Changes in salt bridges following GTP binding in the closed BDLP. **A** The time series of the K154–E438 distance in holo-BDLP is shown (left). A salt bridge is considered to be present when the distance between the N atom of the Lys side chain and either O atom of the Glu side chain is below 0.45 nm (dotted yellow line). Breaking of this salt bridge releases the trunk from the G domain in that area (middle). The motions of the G domain and the stalk are indicated by yellow arrows, and flap1 and flap2 being highlighted in red. The initial and final protein conformations of the MD simulation are shown as light-gray and black cartoon, respectively. Zooming into the K154–E438 region shows how flap1 and the trunk moved away from each other, breaking that salt bridge (right). **B** Occupancies of salt bridges that strengthen (left) or weaken (right) as a result of GTP binding. These strengthened and weakened salt bridges are located at different longitudinal sides of the trunk, giving rise to a shearing motion.

**Figure 7:**
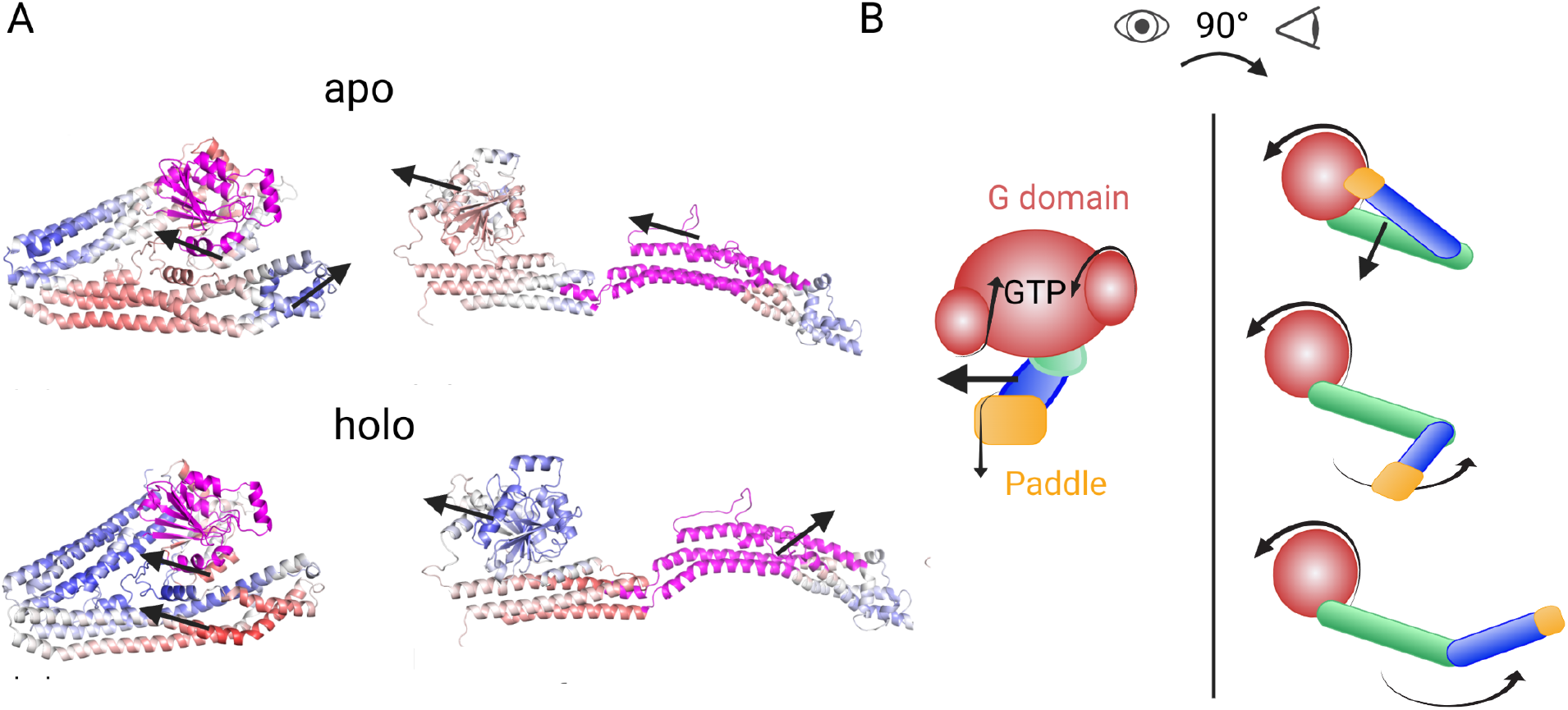
Normal mode analysis of BDLP initial conformations. **A** The normal mode correlation matrix projected onto the structure, where blue indicates anticorrelated and red correlated motions. The regions used as reference are colored in magenta. A reversal of correlations is observed for two cases, when using the flap1-half of the G-domain and the trunk as reference. Arrows are added to indicate the direction of selected motions. More data from that analysis can be found in Figure S2. **B** The principal motions are summarized in a cartoon.

### 2.4 BDLP opening takes less than GTP hydrolysis

Since neither in the standard MD nor in the HREMD simulations the open-to-closed (or closed-to-open) pathway was fully sampled but only led to the semi-closed state, we resorted to USMD to enforce that transition in both apo- and holo-BDLP and calculate the corresponding free energy (also called potential of mean force here^36^). To create initial structures for the USMD windows, we employed a pulling simulation for the closed-to-open transition of apo-BDLP (see Figure S3A for the pulling force applied). That pulling simulation once again confirmed that the hinge1 opening is accompanied by lateral motions of the trunk. Moreover, during the pulling simulation of apo-BDLP, hinge1 opening also coincides with an opening of the GTP binding pocket, an effect which is already visible in Figure 5. For the USMD windows we preferentially used structures sampled in the unbiased simulations and only for the missing parts of the complete pathway, conformations from the pulling simulation were taken. We obtained a range of ca. 100 intermediate initial conformations between *ξ* = 0.8 nm, which is the neck–trunk distance of the closed conformation, and *ξ* = 11 nm for the open state. These windows were restrained with harmonic potentials (see Figure S3B for the distribution of the windows) and simulated for 100 ns. The potential of mean force (PMF) obtained from applying the weighted histogram analysis method (WHAM) to the USMD simulations data of apo-BDLP is shown in Figure 8. We checked for convergence of these simulations by extending each window to 300 ns. The resulting changes in the free energy profile are marginal, and therefore 100 ns is used here (Figure S3C). The complete opening of apo-BDLP requires more than 60 kJ/mol. The initial steep rise of the energy profile between *ξ* = 1 and 4.5 nm can be attributed to the breaking of several salt bridges between the neck and trunk. At *ξ ≈* 5 nm, a metastable conformation corresponding to the semi-closed state already sampled in the MD and HREMD simulation is encountered. This structure is stabilized by a loop of the trunk (490–510) becoming a *β*-hairpin that forms a contact with the G domain. Once this contact is broken, the opening proceeds until an energy valley corresponding to (almost) fully opened apo-BLDP conformations is hit. We repeated the USMD simulations after having docked GTP into the GTP binding site of the starting conformation of each window and adding Mg^2+^ to it. The PMF after 100 ns per US window is shown in Figure 8 (for the error estimate, see Figure S3D). The overall energetic threshold for the closed-to-open transition is lowered to about 30 kJ/mol. Thus, the binding of GTP renders the open conformation more favorable.

**Figure 8:**
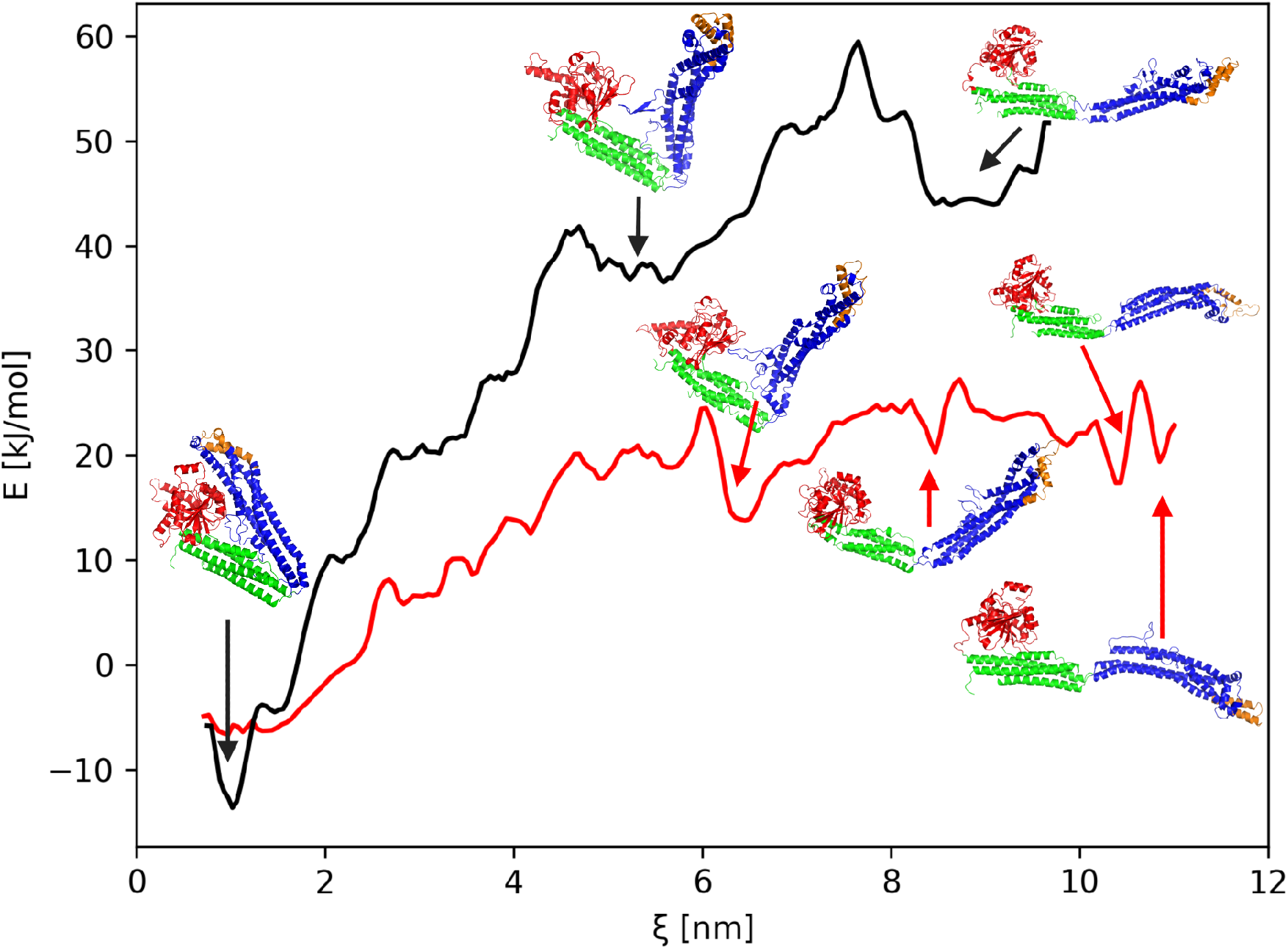
The free energy of the close-to-open transition of BDLP. These results are obtained from umbrella sampling MD simulations apo-BDLP (black) and holo-BDLP (red). The energies smoothed with a uniform filter. Representative structures of local minima are show.

As discussed above, the addition of GTP sets off a number of structural changes in the G domain, which in turn break key salt bridges between the G domain and the trunk, enabling the opening at hinge1. Hoewever, also hinge2 is affected by GTP binding, as it causes the G domain to roll away from the neck, which lowers the energy barrier for dissolving the interaction between the loop of the trunk and the G domain that is present in the intermediate at *ξ ≈* 6.5 nm. It should be noted, however, that both 60 kJ/mol and 30 kJ/mol are in principle accessible by the hydrolysis of a single GTP molecule,^37,38^ which releases 60 kJ/mol.

On the other hand, one also needs to consider the time scales involved for the different energetic barriers. Using the Arrhenius equation, *τ* = *τ*_0_ exp(Δ*G*^#^*/k*_B_*T*) with Δ*G*^#^ as the overall free energy barrier, *T* = 310 K, *k*_B_ being the Boltzmann constant, and *τ*_0_ *≈* 10^*−*12^ s at 310 K, we obtain that, on average, more than 7 ms are needed to for the closed-to-open transition of apo-BDLP, while this reduces to 100–200 ns for holo-BDLP. The conformational change of BDLP can thus take place more quickly than GTP hydrolysis and before the dimer dissociates, and is well below the seconds to minutes reported for membrane fission itself.^17^ A valid question though is why we did not sample that transition in the 600 ns MD or HREMD simulation of holo-BDLP. The most likely answer is that with USMD one concentrates on a narrow part of the energy landscape between two end states, while in unbiased simulations the protein has more possibilities to explore its conformational space and does not necessarily strike out to the end state we expect it to develop to. The same observation we made in a study of a much smaller movement, of a loop in triosephosphate isomerase that switches between an open and closed conformation depending on the substrate-loading state.^39^

## 3 Discussion

Combining the analysis of the BDLP conformations and that of salt bridges with the knowledge gained about the forces needed for the closed-to-open transition, we deduce that the energetic barrier for BDLP opening mostly consists of the breaking of six salt bridges between the neck and trunk, after which less force is needed to complete the opening. The closed-to-open transition not only involves a hinge1 movement, but involves a shearing motion in lateral direction to minimize the electrostatic force needed. The open form of BDLP is accessible by the energy released from the hydrolysis of a single GTP molecule, and just the binding of GTP lowers the energy barrier for the opening. This allosteric action of GTP binding is relayed from the GTP binding site via the motions of two flaps in the G domain, closing to cover the GTP-loaded site, which breaks salt bridges between the G domain and the trunk and initiates the detachment of the paddle from the G domain. Salt bridges on the other longitudinal side of the trunk are formed, which guides the lateral motion of the stalk region during opening, while the release of salt bridges between stalk and G domain cause the latter to roll away at hinge2 (Figure S1). However, further hinge1 opening coincides with an opening of the GTP binding pocket (Figure 5), which explains why GTP hydrolysis by dynamins usually requires the dynamins to be polymerized where the G domains would stabilize each other to provide the structural stability needed for the hydrolysis reaction to take place. The mechanism of binding GTP and hinge1 opening must thus be connected in both directions, since the open form is more likely to be assumed with GTP (and not GDP) bound, in polymeric, membrane-associated BDLP.^22^ Vice versa, the polymeric form must communicate its status to the GTPase domain, in order to stimulate GTPase activity, and possibly to facilitate GDP release afterwards. We thus agree with^21^ that GTP binding causes the hinge1 opening and provide the mechanistics behind that transition. The PMF of the open-to-closed transition in Figure 8 further shows that the open state of the BDLP monomer in solution is less stable than than the closed state, which clarifies why Low et al. only observed the open holo-BDLP form when being membrane-bound and polymeric. Importantly, GTP hydrolysis cannot be the cause for the opening, since membrane tubulation also happens with non-hydrolyzable GTP analogues. Moreover, GDP is thought to cause depolymerization and membrane dissociation.^22^ The insights gained about BDLP’s mechanism of action may be transferable to other DLPs, contributing to our understanding of fundamental cellular processes, such as endocytosis, cell division, and immune response.

## 4 Methods

The open and closed structures of BDLP were taken from the Protein Data Bank (2J68 and 2W6D)^22,23^ and completed using RCD+, GalaxyLoop, DaReUs-Loop and ModLoop.^40–43^ In the all simulations, AMBERff99SB*ILDNP^44^ and the TIP3P water model were used for modeling the protein and its surrounding. After energy minimization, MD simulations for equilibration in the NVT and NpT ensemble were performed, followed by production MD runs, HREMD or USMD. The protein was sampled in its apo form (4 fs timestep)and with GTP and Mg^2+^ bound (2 fs timestep), called apo- and holo-BDLP henceforth. Depending on the protein conformation, the system sizes were between 300,000 and 1,000,000 atoms. All simulations, including the HREMD and USMD simulations, were performed with GROMACS (version 2016.4).^26^ For the HREMD simulations, 30 replicas with 100 ns per replica were used, resulting in an average acceptance rate of 30% for exchanges between the replicas. For the USMD simulations, 74 and 67 windows with 100 ns per window were used for apo-BDLP and holo-BDLP, respectively. To test for convergence, the USMD windows for apo-BDLP were extended to 300 ns. Initial conformations for the windows were generated in preceding pulling simulations, and the resulting conformations were restrained with a harmonic potential in the USMD simulations. The free energy profile from the USMD simulations was obtained from a WHAM analysis. ^36,45^ All simulations performed in this study are summarized in Table S1; they yielded an accumulated simulation time of 19.5 *µ*s. Different GROMACS tools were used for analysis, among them the calculation of the RMSF, various distances and angles, spatial distributions, cluster analysis, and principal component analysis (PCA). The RMSF values between 0.1 and 4 Å were projected onto the structures, with red areas denoting flexible residues and blue areas rigid ones (RMSF *>* 2 Åand ≥ 2 Å, respectively). The distances and angles used to characterize the hinge movements are defined in Figure 1. To calculate the displacement of the stalk tip compared to the initial position, denoted as Δ*x*, Δ*y*, and Δ*z*, a custom Tcl script was used in VMD^46^ text mode. Here, Δ*z* corresponds to lateral motions of the trunk. The VMD-integrated tool ProDy^47^ was used to study normal modes of the initial structures. Free energy surfaces were created using a custom Python script. In the analysis of the HREMD simulations, only the corresponding target replica was used. Figures were generated with PyMOL, Inkscape, and BioRender.

More details about the simulation and analysis methods are provided in the Supplementary Information.

## 5 Acknowledgments

B.S. gratefully acknowledges funding from the Deutsche Forschungsgemeinschaft (DFG, German Research Foundation) (project A07 of the CRC 1208). W.S. is grateful for a PhD fellowship from Jürgen Manchot Stiftung. The project was made possible thanks to computing time granted through the Gauss Centre for Supercomputing e.V. (www.gauss-centre.eu) on the GCS Supercomputer SuperMUC-NG at Leibniz Supercomputing Centre (www.lrz.de) (project pn98zo).

## 6 Author contributions

B.S. designed the research. W.S. performed the simulations and data analysis. W.S. and B.S. wrote and reviewed the manuscript. B.S. supervised the project.

## 7 Additional information

Supplementary information is available.

## Notes

### Competing Interest Statement

The authors have declared no competing interest.

