## Supplementary Methods, Tables and Figures for "Allosteric communication induced by GTP binding sets off a closed-to-open transition in a bacterial dynamin-like protein"

### 1 Supplementary methods

The structures of BDLP were taken from the Protein Data Bank (2J68 and 2W6D)[1,2] and modeled using RCD+, GalaxyLoop, DaReUs-Loop and ModLoop[3–6]
For the MD simulations, the Amber99SB*-ILDNP force field(,[7–9] TIP3P water[10] and GROMACS 2016[11] were used, at a temperature of 303 K (37°C) and a pressure of 1 bar. In Table S1, all MD simulations performed in this study are summarized, which corresponds to a total simulated time of 19.5 $\mu$s. GTP was parametrized according to the protocol described in[12] and docked to BDLP using AutoDock.[13] Several unrestrained simulation of BDLP were performed in order to explore the conformational space. The system sizes are listed in Table S1. 8 or 10 $Na^+$ were added in order to neutralize the apo- or holo-protein's charges, respectively, with an additional $Mg^{2+}$ being added to the holo simulations. First, an energy minimization was performed using a steepest descent algorithm, followed by a subsequent 0.1 ns NVT equilibration and a 1 ns NpT equilibration. During these steps, the protein's heavy atoms were restrained with a force constant of 10 kJ/mol*Å$^2$ . The velocity rescaling thermostat was employed to regulate the temperature in the NVT simulations, while the Nosé-Hoover thermostat[14,15] and the isotropic Parrinello-Rahman barostat[16] were used for the NpT simulations. The particle mesh-Ewald (PME)[17,18] sum method was used to calculate electrostatic interactions with xyz-periodic boundary conditions. The van der Waals (vdW) and short-range Coulombic interaction cutoffs were set to 12 Å, repulsive Lennard-Jones (LJ) interactions were cut at 10 Å. The equations of motion were integrated using the leapfrog method, and the LINCS algorithm[19] was used to constrain all bonds. For the production rund, the same setup as in the NpT equilibration was used, minus the position restraints.

In the apo-BDLP systems, some hydrogen atoms were treated as virtual interaction sites, permitting an integration time step of 4 fs while maintaining energy conservation.[20] For the holo-BDLP systems, a time step of 2 fs was used. Coordinates and velocities were recorded every 20 ps in both cases. In all simulations, we applied position restraints for the rigid $C_\alpha$ atoms of the G-domain $\beta$-sheets to remove overall translation and rotation, allowing us to decrease the box size without harming the protein's flexibilities.[21] To ensure that GTP stayed in its binding pocket, two distance restraints between GTP-atoms O3G/O2S and Ser97/Val243 of BDLP were applied using the GROMACS pull code, when neccessary.

Next, we performed a Hamiltonian replica exchange MD (HREMD) simulation[22] with 30 replicas, each 100 ns per replica in length. The energy function of BDLP, and its protein-water interactions were modified in each but the target replica by applying biasing factors of 310 K/$T$, with the 30 temperatures $T$ exponentially distributed between 310 and 370 K ($1 < \lambda < 0.667$). The unbiased target replica at 310 K was used for analysis. The average exchange probability between the replicas was ca. 30%. The HREMD simulations were conducted with Gromacs 2016.4 patched with the PLUMED plugin (version 2.4.1) ([23]). In all HREMD simulations, we used the v-rescale thermostat with canonical sampling and the Parrinello-Rahman barostat.[16] Apart from that, the setup was identical to the regular production runs described above.

**Table S1:** Simulations performed in this study, amounting to a total of 19.5 $\mu$s.

| simulation type | conformation | apo/holo | system size (atoms) | length |
|---|---|---|---|---|
| MD unbiased | open | apo | 238,657 | 600 ns |
| MD unbiased | open | holo | 194,959 | 600 ns |
| MD unbiased | closed | apo | 284,578 | 600 ns |
| MD unbiased | closed | holo | 194,905 | 600 ns |
| HREMD | open | apo | 238,657 | 30 × 100 ns |
| MD pulling | closed-to-open | apo | 238,657 | 0.8 ns |
| USMD | – | apo | 238,519 to 1,537,576 | 74 × 100 ns |
| USMD | – | holo | 787,203 to 1,536,653 | 67 × 100 ns |

From the unrestrained simulations, the predominant motion was identified using principal component analysis. This motion was then described using a distance, measured between the $C_\alpha$ atoms of residues 224 and 453. This order parameter/reaction coordinate is referred to as $\xi$ and used for the USMD simulations. In the USMD, 100 apo-BDLP conformations along $\xi$ were simulated for 100 ns each. The force constants for restraining the conformations along $\xi$ ranged between 3 and 6,200 kJ/(mol·nm$^2$), depending on the stability of the different windows; though the majority of the windows were simulated with 500 kJ/(mol·nm$^2$). This setup was repeated for holo-BDLP. Of the 100 windows, only those which stayed in the range of the original window, and those where GTP remained bound at least 50% of the time were used for analysis.

The analysis was mainly performed with GROMACS-internal tools and included following calculations:

**WHAM** WHAM[24] was used as implemented in GROMACS,[11] for 74 (apo) or 67 (holo) trajectories of 100 ns each, at a temperature of 303 K. To ensure convergence, a subset of windows was extended to 300 ns, however, the energy profile did not change anymore after 100 ns. The profile shown in the main text was smoothed with a window size of 5 in reflect mode, while the extended data shows the original profile. Error bars were generated by 1000-fold bootstrapping, as implemented in *gmx wham*.

**Distances** Distance $\xi$ was measured between the $C_\alpha$ atoms of residues 224 and 453.

**Angles** Angle $\alpha$ defining hinge1 was measured between the $C_\alpha$ atoms of residues 4, 359, and 587. Angle $\beta$ defining hinge2 was measured between the $C_\alpha$ atoms of residues 291, 303 and 323.

**RMSF** The *gmx RMSF* tool was used to calculate fluctuations of the $C_\alpha$ atoms around their time-averaged positions. The resulting values (between 0.1 and 4 Angstrom) were projected onto the structue, with red areas denoting flexible residues (>2 Å) and blue areas rigid ones (<2 Å).

**Spatial distribution** In order to visualize the area sampled by the stalk tip, the *gmx spatial* tool was used to calculate its spatial distribution. First, the trajectory was fitted to the G domain of BDLP, then the position of the $C_\alpha$ atom of residue 580 was binned at 0.1 nm precision. The resulting distribution was visualized in PyMOL.

**PCA** To observe the largest motions present in the trajectory, principal component analysis of the atomic motions was performed, using the PCA implementation of GROMACS. The trajectory was fitted to the backbone for this analysis. The three main backbone motions were visualized both for the whole protein and the flexible parts of the G domain (flap1, flap2, hinge2, switch2).

**Cluster analysis** For the clustering of the G domain motifs, the trajectory was first fitted to the $C_\alpha$ atoms of the G domain with a time step of 60 ps between snapshots. The clustering was then performed only on the motifs (flap1, flap2, switch1, hinge2, $Mg^{2+}$, and GTP) with a cutoff 0.25 nm and the *nofit* option.

**Normal mode analysis** The VMD plugin ProDy[25] was used to analyze the normal modes of the initial open and closed BDLP conformations, both in their apo and holo forms. The anisotropic network model was calculated for the $C_\alpha$ atoms and extended to the backbone, with a cutoff of 19 nm.

#### 2 Supplementary figures

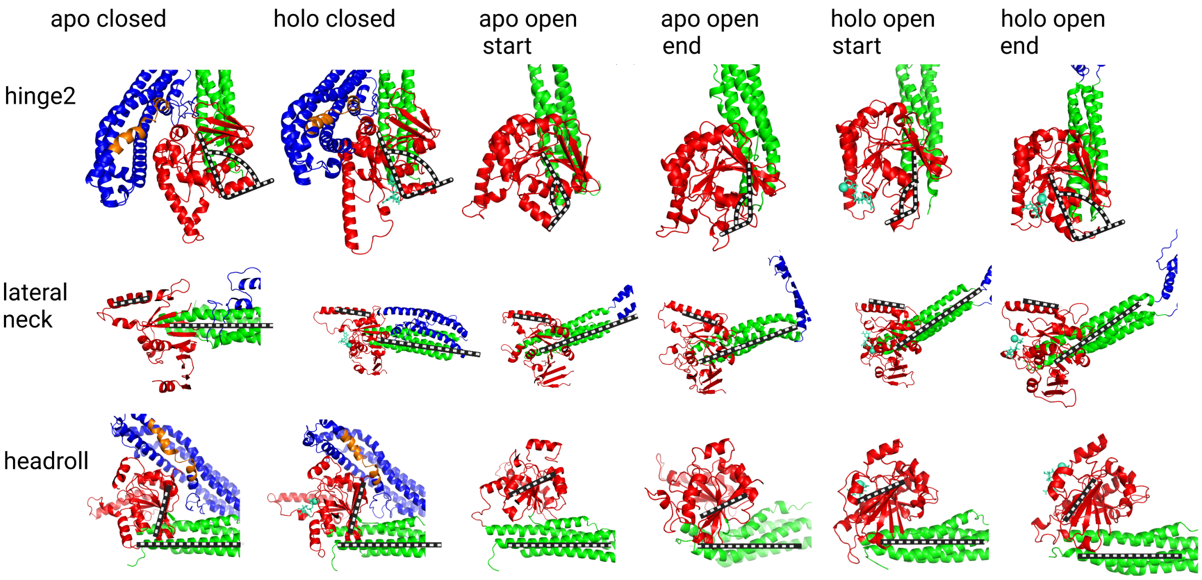


**Figure S1: Snapshots of different hinge2 motions.** Hinge2 is defined by the angle indicated in the top row. A more structured hinge2 region leads to a 90°angle between helix12 and helix13. The lateral motion of the stalk beginning at helix13 becomes apparent in the second row, when using helix10 as a reference. In the third row, we can see how $\beta$-strand4 is at a 90°angle to the stalk in the closed form, but more parallel for the open form. These differences could be largely originate from the initial starting structures and are equilibrated after the simulations.

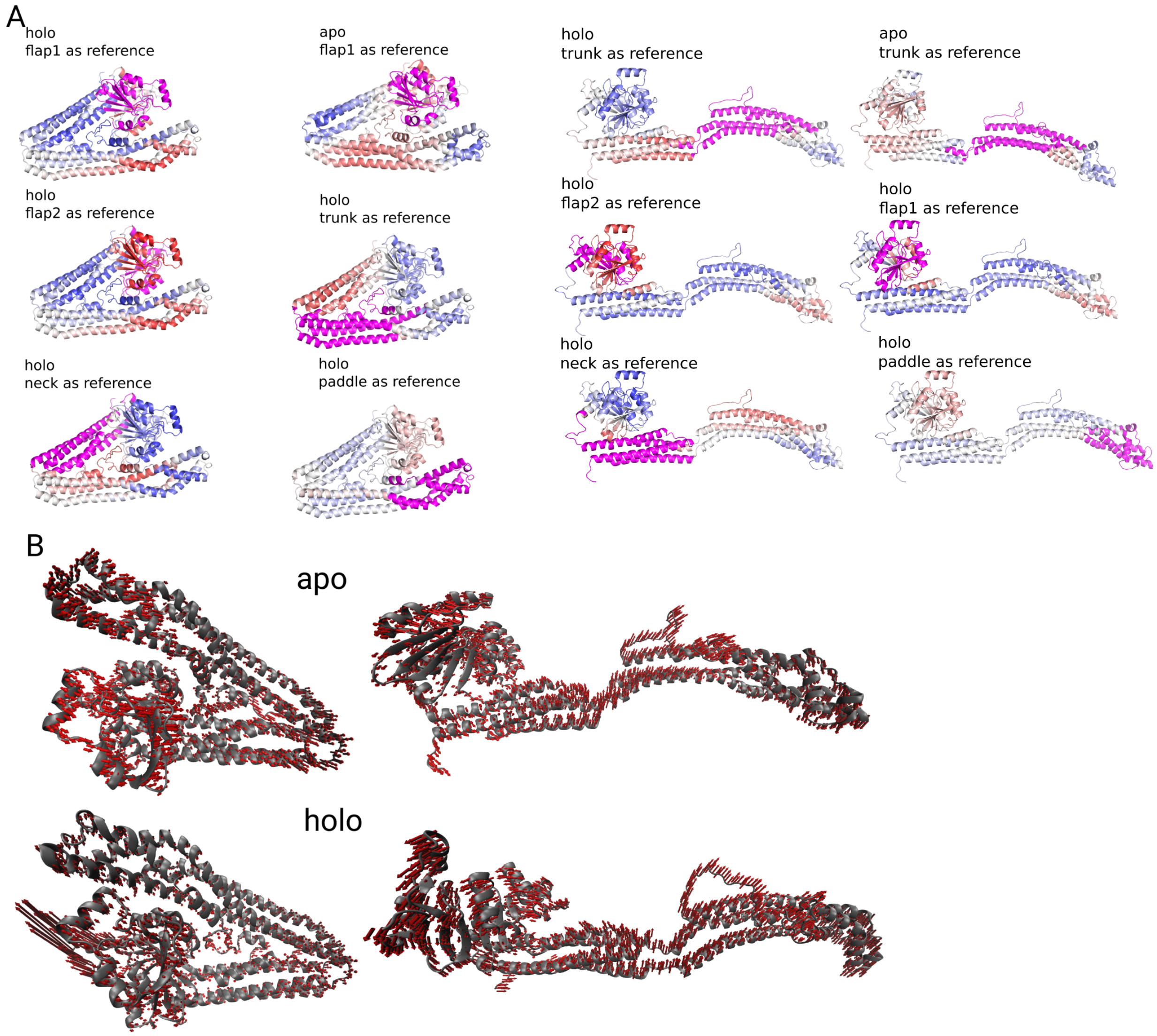


**Figure S2: Normal mode analysis of the BDLP starting conformations. A** The first normal mode projected onto the structure as red arrows. **B** The correlation matrix projected onto the structures (blue = anticorrelated motions, red = correlated motion, magenta = reference region). The correlations in the apo state are only shown for one reference region, to illustrate the most striking differences to the holo state.

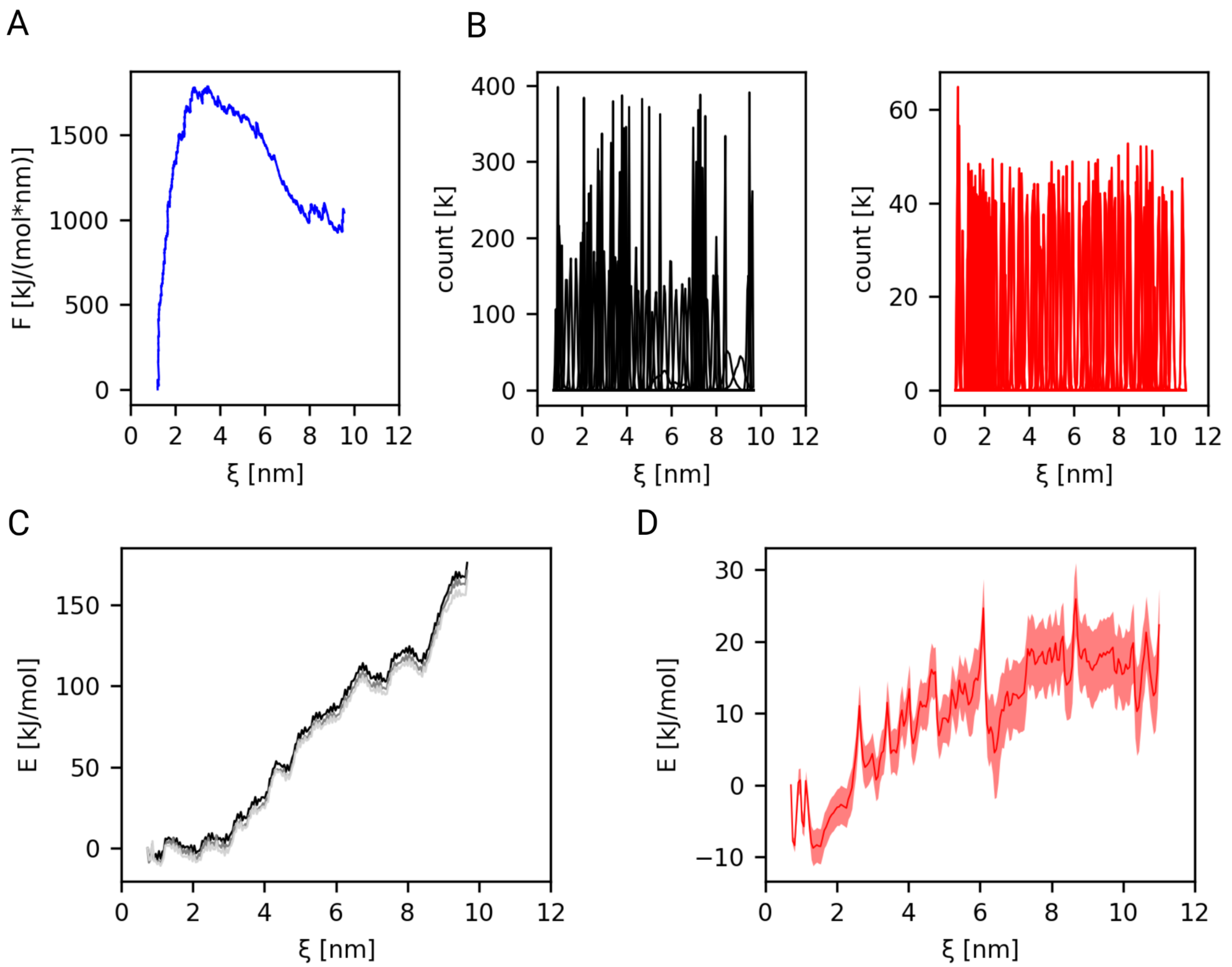

**Figure S3: Umbrella sampling MD simulations of BDLP. A** Force profile during the initial pulling simulation. **B** Coverage of the order parameter $\xi$ by the windows in the USMD simulations of apo-BDLP (left) and holo-BDLP (right). **C** Convergence test for the USMD simulation of apo-BDLP using increasing simulation lengths: 100 ns (black), 200 ns (grey), 300 ns (light grey). **D** Margin of error for the USMD simulation of holo-BDLP, generated with 1000-fold bootstrapping.

(3) Park, H.; Lee, G. R.; Heo, L.; Seok, C. Protein loop modeling using a new hybrid energy

function and its application to modeling in inaccurate structural environments. *PLoS ONE* **2014**, *9*.
